# Evaluation of the capacity of Pediatricians in Nigeria to conduct of research: A nationwide Survey

**DOI:** 10.1101/348128

**Authors:** Maduka D. Ughasoro, Iliya Jalo, Angela Okolo, Ebun Adejiyugbe, Mariya Murktah, Austin Omogberare, Ben Onankpa, Ngozi Ibeziako, Agozie Ubesie, Damian Nwaneri, Clement Ezechukwu, Oguche Stephen

## Abstract

**Background:** Quality health care service delivery to children and adolescents is enhanced by continuous research into the health challenges of this subpopulation led by paediatricians with tremendous capacity to investigate and proffer solutions to the myriads of childhood illnesses. Understanding the health issues therefore is the foundation for implementation of viable interventions that assure optimum service delivery. In view of this background, the Paediatric Association of Nigeria (PAN) directed that research into children’s health challenges in Nigeria should be brought to the front burner in the country. Pursuant to this laudable goal this study was conceived to evaluate the research capacity and capability of paediatricians in Nigeria and the institutions they represent. In view of above needs, this study aimed at evaluating the research capacities and challenges among paediatricians.

**Methods:** The survey used a cross-sectional nationwide design to enroll paediatricians into the study. The study was a combination of both online and face-to-face survey using questionnaire developed from Research Capacity Assessment Framework. Information on previous research work, challenges encountered, existing capacity and utilization of research outcome were obtained. The SPSS version 20 was used for data entry and analysis. For qualitative variables, similar responses were grouped under thematic heading.

**Results:** The response rates for online (via email survey, group-administered in a conference and individual face-to-face (at workshops) were 32 (3.2%), 75 (13.6%) and 15 (60%) respectively. The majority, 87(85.5%) of the participants had conducted prevalence studies, compared to 9 (8.8%) that had done experimental studies. Those who have ever received grant funding for their studies were 21 (19.4%), while the proportion whose research outcome had informed policy update and practice were policy 20 (18.2%). More than 55% of the participants had challenges on some of the seven aspects of research: research topic, proposal, funding, fieldwork, analysis, utilizing findings and collaboration. Less than 40% of the participants had received training on some of the tested 14 research capacity areas except for the area of ethics where 78 (70.9%) reported having received training. For 51 (46.4%) this ethics training included the Good Clinical Practice Guidelines.

**Conclusion:** Nigerian Academic Paediatricians need to be stimulated to develop interest in research by building their presently low research capacity if future paediatric practice is to be driven significantly by evidence.

## Introduction

It is the responsibility of paediatricians among other child advocates and stakeholders around the world to ensure optimal wellbeing of every child. This is to be achieved through their involvement in these five inter-related domains of practice: clinical service and consultancy, research, education/teaching, clinical leadership and clinical service planning and management.^1^ The benefits to child healthcare service delivery can be improved if human resources are allocated effectively to cover all these domains. However, in reality, this does not appear to be the case. One domain that suffers the most is clinical research. The fact remains that the entire deliverables in paediatric care can be greatly enhanced if informed by evidence-based approaches. Paediatricians are exceptionally well positioned to lead in this effort because of both their specialization and positioning to utilize these evidenced-based clinical practice guidelines to improve child health care. This is possible if the health research is designed to address key clinical needs and resolve knowledge gaps in order to provide practical and implementable solutions. ^2^ Clinical research is fundamental in health care delivery system. When the research is carried out in a goal-oriented manner with clear view of expected outcome, is expected to propel continuous improvement in the disposition of healthcare workers to towards service delivery. Such efforts should be rewarded especially where the outcomes has significant impact on clinical practice by closing knowledge gaps. ^2^

Although several studies have cited lack of support systems for research, increased clinical workload,^3^ lack of funds, and the high premium placed on clinical service and consultancy as reasons for paucity of clinical research activities, ^4^ the extent of involvement of paediatricians in clinical research has not been well documented. Research into childhood illnesses and consequent publications abound in almost all aspects of specializations. However, what has not been fully executed is the development of prioritized research agenda for Nigeria. Agenda setting will focus on critical child and adolescent health needs. This led the Paediatric Association of Nigeria (PAN) to inaugurate a Research Committee (PAN-RC) in April 2017 charged to strengthen and harmonize research activities among paediatricians, contribute and complement child health research efforts in the country and advance research knowledge and skills of its members. The committee embarked on a stepwise approach to develop a child and adolescent health research priority list for researchers and other stakeholders interested in finding solutions to childhood diseases in Nigeria. The first step in the process is a need assessment to determine the existing and spread of research capacities among paediatricians in Nigeria. Using a SWOT analysis (strength, weakness, opportunities and threat) approach based on the research capacity assessment framework the study aims to evaluate the paediatricians’ clinical research engagements, challenges, and capacities to undertake meaningful research. This paper reports the preliminary findings on research capacities of paediatricians in a nationwide survey.

## Material and Methods

### The Setting

Paediatrics Association of Nigeria (PAN) is the umbrella association of all academic and practicing paediatricians in Nigeria with over 1000 members. Its members provide healthcare to children and adolescents in healthcare facilities all over the country including but not limited to training of undergraduate medical students and resident doctors in addition to providing consultancy services to national and international governmental and non-governmental organizations. The association aspires to see that child and adolescent health interventions are based on local evidence in addition to global observations.

### Study Design

The study used a cross-sectional questionnaire based survey. A combination of both online and physical survey utilizing both quantitative and qualitative research methods was used.

## Development of Study Tool

A review of available literature on assessment of research capacity of healthcare workers was conducted. The findings of the systemic review informed the development of the questionnaire. The survey questionnaire is based on the research capacity assessment scoring framework developed by Dana et al ^5^ designed to explore competence of researchers to conduct of clinical research. The survey questionnaire has some close and open ended fields about the age, gender, place of work, position in service (cadre), institutional affiliation, area of expertise, participation in a research work as principal investigator, the type of research work, what influenced the choice of research area/topic, challenges/barriers experienced conducting the research, if previous research findings had influenced evidence-based change in policy or practice, how the policy/practice change was achieved, if ever received grants, source of such grant, purpose for the grant and if aspects of health research pursued had received training,

## Online survey

The paedaitricians were invited by email using the Paediatric Association of Nigeria (PAN) mailing list to participate in the online survey.^6^ The informed consent form was administered to study participants online inviting them to participate in the study. Online lasted over 6 weeks and the there was weekly reminder. The participants who returned their completed consent forms accepting to be part of the survey received the survey questionnaire. Questionnaire distribution was extended to paediatricians practicing in the six geopolitical regions of the country, viz North West, North Central, North East, South West, South East and South-South regions.

## Structured interviews

Paediatricians who attended Training of Trainers (TOT) workshop on Management of Severe Acute Malnutrition (SAM) in Asaba August 2017 and the annual Paediatric Association of Nigeria Conference (PANConf) in January 2018 where invited to participate in the survey. Those who had filled the online survey were excluded from participating at this level. The interview questionnaire was the same sequence as the online survey tool. The open ended questions enabled gathering indepth information on what influenced the choice of research area/topic and how research translated to policy and practice. To avoid bias and influencing the responses given by the participants, interviewee-administered (self-administered) method was used instead of interviewer-administered method. A total of 25 questionnaires were distributed during the TOT on SAM workshop in Asaba and 15 questionnaires were returned completed. During the PANConf 2018, a total of 550 questionnaires were distributed, and 75 completed questionnaires were returned. To increase the number of completed questionnaires, additional distribution of survey questionnaires was conducted in the six geopolitical zones of the country through contact persons. This yielded 17 more completed questionnaires. A total of 117 completed questionnaires were received. The scores for the level of challenges were categorized under even Likert scale of no challenge (0), low challenge (1), medium challenge (2.4) and high challenge (5 – 7). Their responses to the open ended questions: what influenced your choice of research area/topic and how policy change was achieved, were grouped under thematic headings.

## Ethical Consideration

The Health Research Ethics Committee of University of Nigeria Teaching Hospital (UNTH) reviewed the research protocol and approved the study. Written informed consent was obtained from the participants before participating in the study.

## Data analysis

The data was double entered into SPSS version 20 software. Frequencies were calculated for discrete variables, while means and standard deviation were calculated for continuous variables. The qualitative data was reviewed and categorized under thematic headings.

## Results

The Questionnaires retrieval rate for online, conference, and workshop were 32 (3.2%), 75 (13.6%) and 15 (60%). The extra 17 questionnaires were obtained from regional survey. A total of 117 questionnaires were obtained, 7 incomplete questionnaires were excluded and 110 were finally analyzed (Figure1).

**Figure 1:**
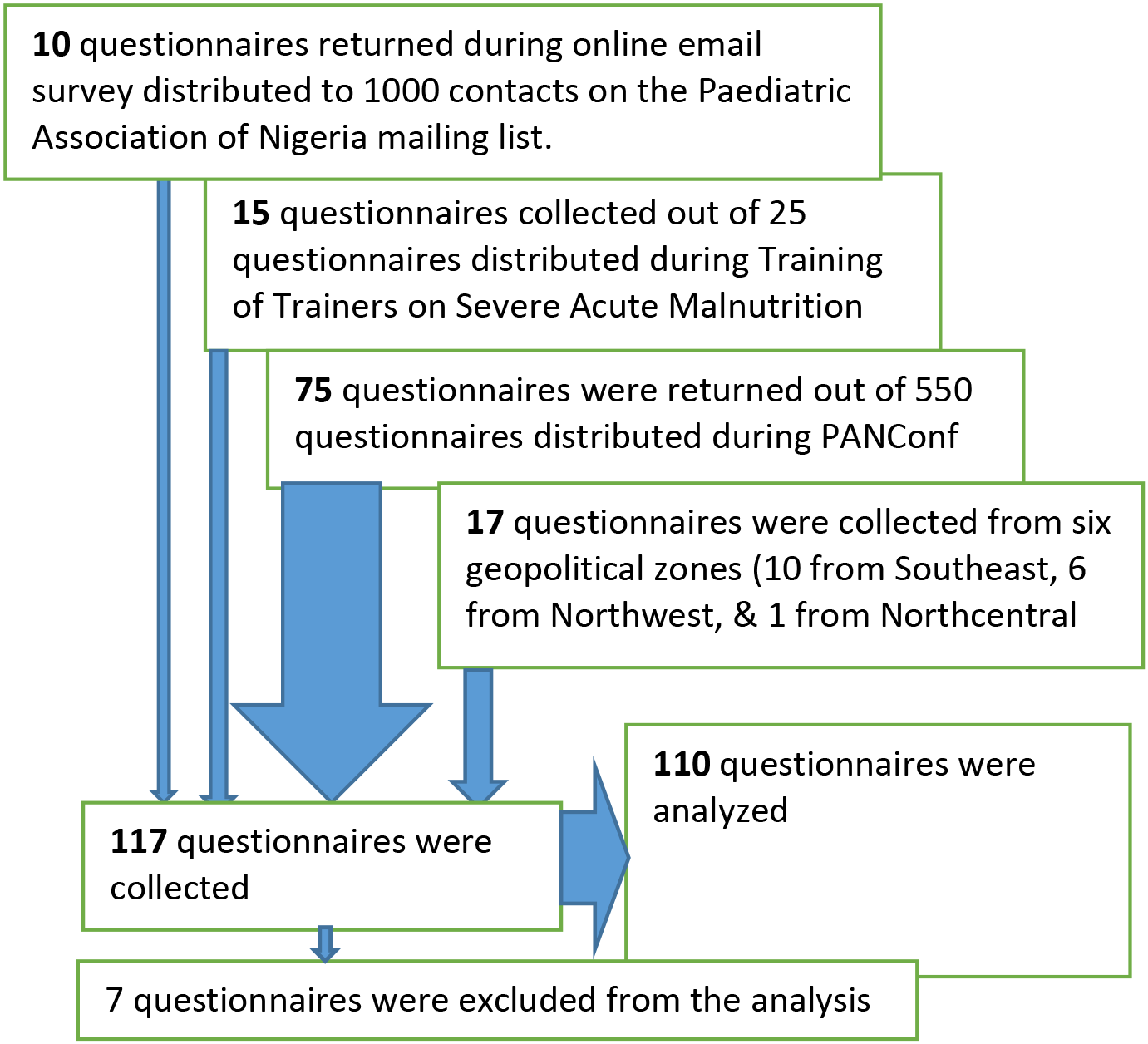
Flow chart of participants.

The demographic characteristics of study participants are shown in table 1. The participants included 40 (36.4%) males and 70 (63.6%) females. The mean age of participants was 44.8 years. Thirty (28.6%) of the participants representing the majority were general paediatricians (Figure 2). Responses from the regions are shown in figure 3.

**Table 1:**
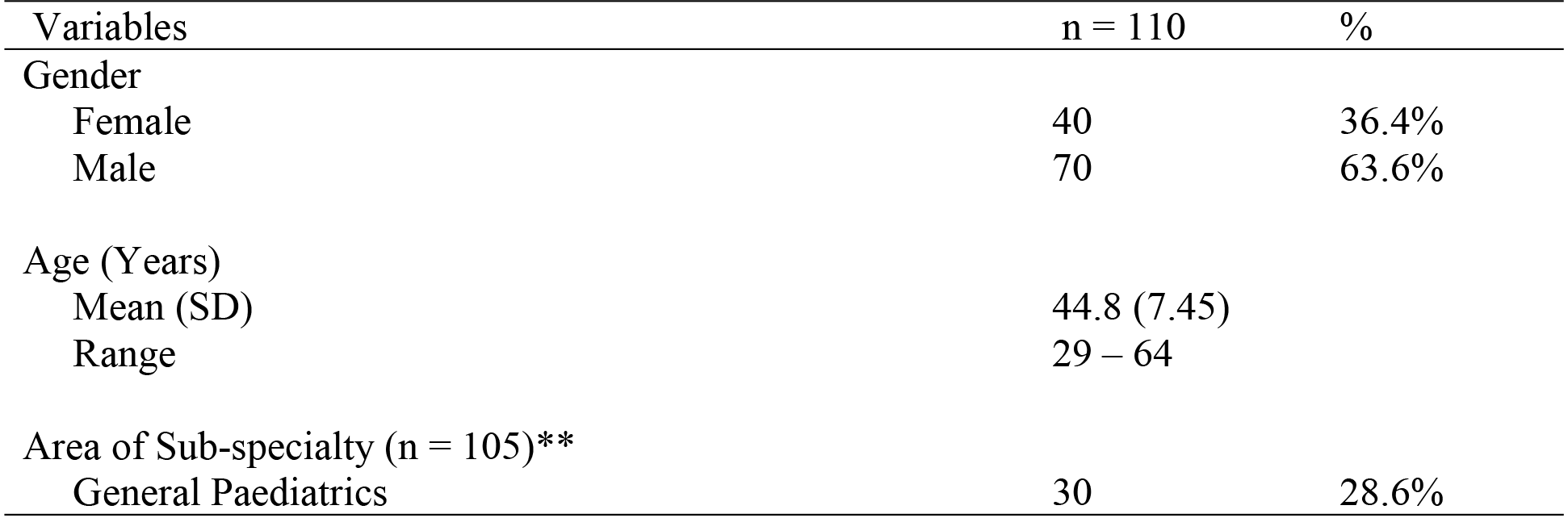
Demographic characteristics of participants

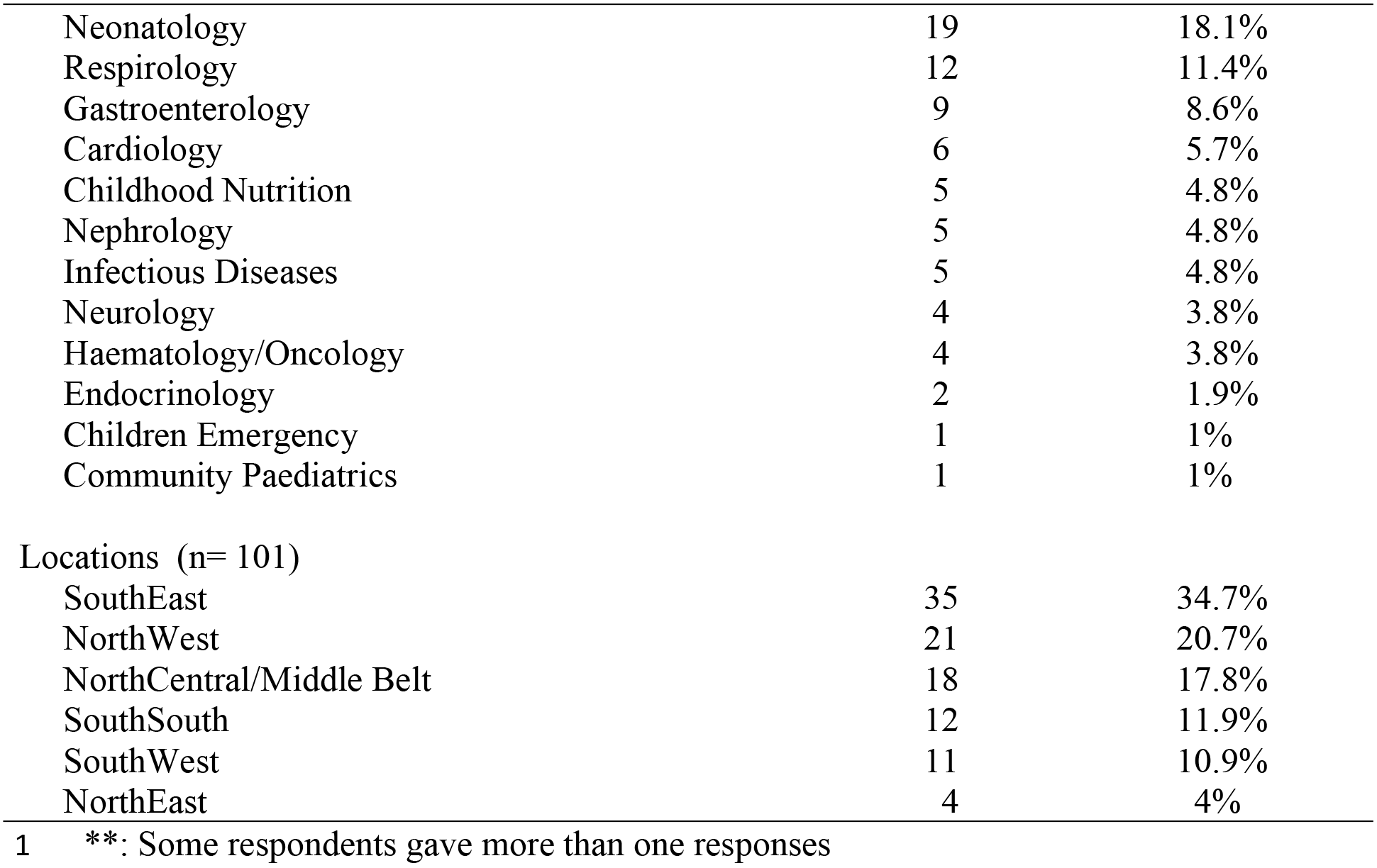

**Figure 2.**
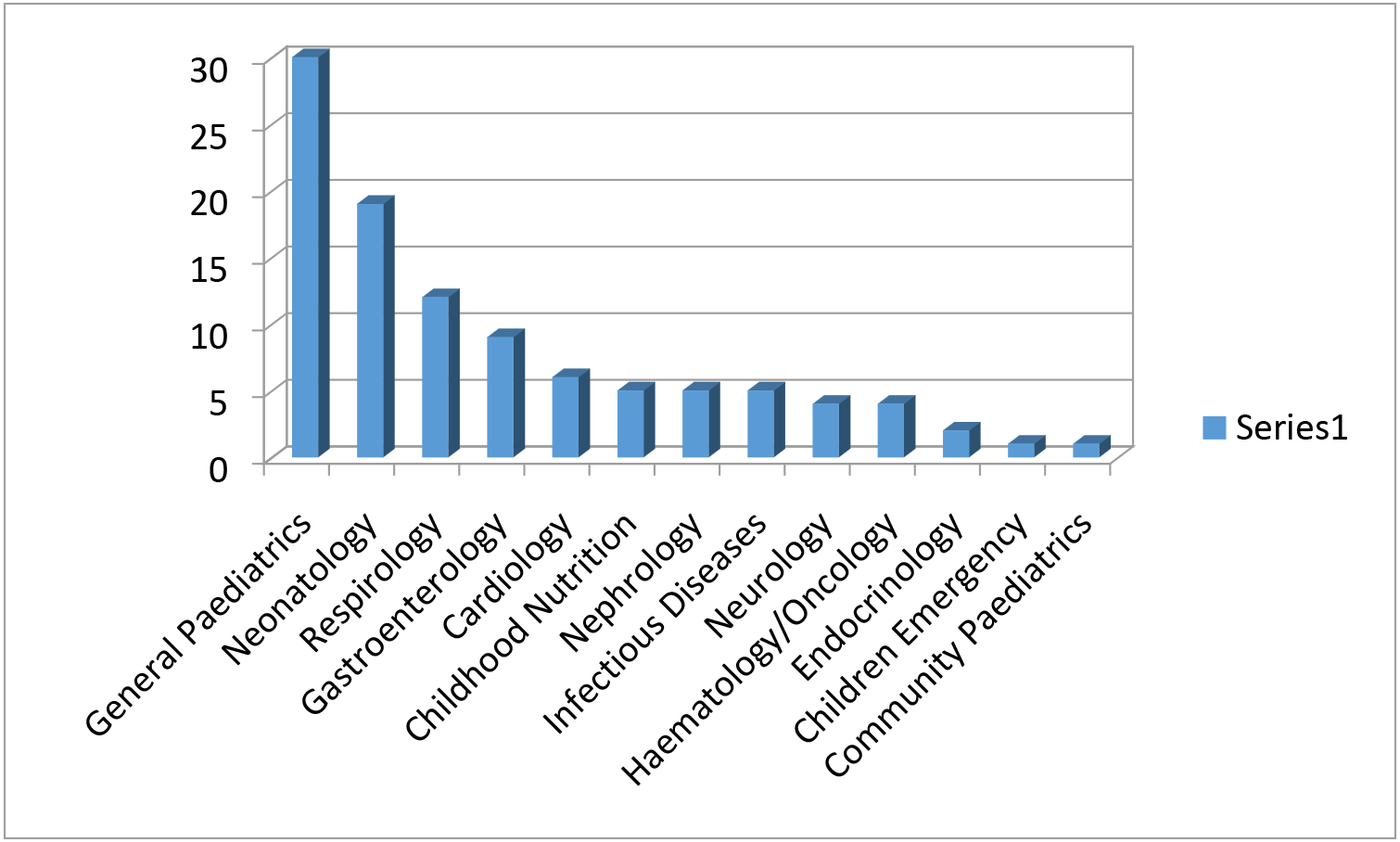
Distribution of Participants according to sub-specialties

**Figure 3:**
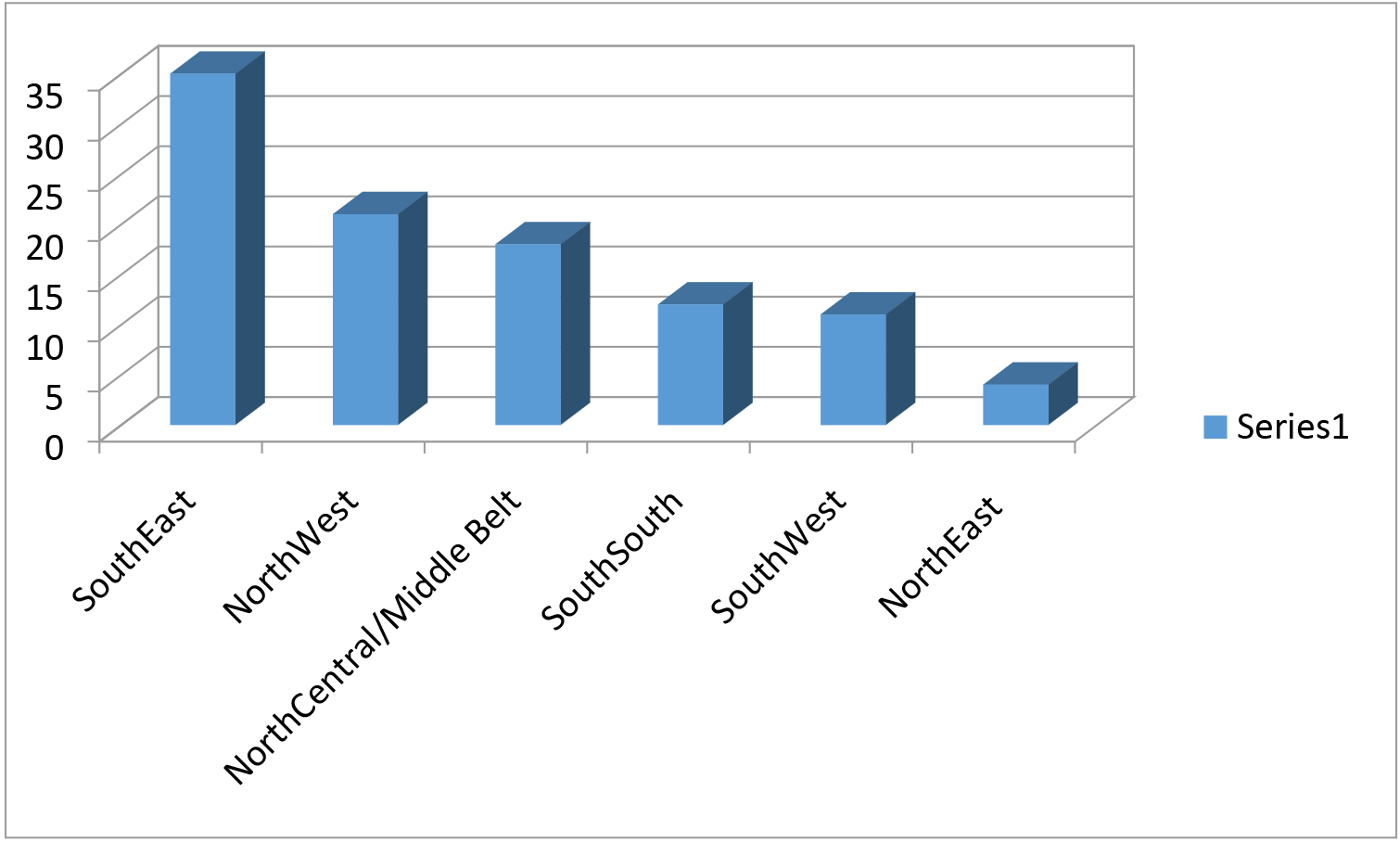
Distribution of participants according to their geo-locations

The number (%) of the participants that had acted as principal investigator in a research was 102 (92.7%). Most, 87 (85.5%), of their previous research works were prevalence studies. The least frequent conducted research was randomized controlled clinical trial, representing 9 (8.8%). The reasons for embarking on clinical research work included: to answer prevailing question/fill an existing knowledge gap 42 (41.2%), found the research work interesting 41 (40.1%), and benefits to patients/clinical observations 33 (32.4%). More uncommon reasons given included: suggestion by funder/collaboration 2 (1.9%) and prompted by rare a condition 1 (0.9%). The majority (80.9%) of paediatricians had not received grants/sponsorship to conduct research. Most of the outcomes of their previous research works had not informed change in policy update or practice. See Table 2 for complete details.

**Table 2:** Scenario of current researches and outcomes

| Variables | n= 110 | % |
| --- | --- | --- |
| Have you ever been a Principal Investigator (PI) in a research? |  |  |
| Yes | 102 | 92.7% |
| No | 0 | 0 |
| Missing response | 8 | 7.3% |
| Which type of research were you the PI? (n=102)** |  |  |
| Prevalence Research | 87 | 85.5% |
| Observational Research | 20 | 19.6% |
| Operational Research | 13 | 12.7% |
| Experimental (Randomized Clinical Trial) | 9 | 8.8% |
| Others (Reviews) | 1 | 0.9% |
| Determinants of research area (n=102)‡ |  |  |
| Answer prevailing question/Existing Knowledge gap | 42 | 41.2% |
| Interest in the area/topic | 41 | 40.1% |
| Benefit to patients/Clinical Observation | 33 | 32.4% |
| Feasibility of conducting the research | 24 | 23.5% |
| Availability of fund | 17 | 16.7% |
| Disease burden (mortality/morbidity) | 13 | 12.7% |
| Needs assessment | 11 | 10.8% |
| Dissertation/Academic appraisal | 9 | 8.8% |
| Suggested by funder/Collaboration | 2 | 1.9% |
| Rare condition | 1 | 0.9% |
| Have you ever received grants to conduct a research (n=108) |  |  |
| Yes | 21 | 19.4% |
| No | 87 | 80.6% |
| Had any of your previous research informed policy change (n = 110) |  |  |
| Yes | 20 | 18.2% |
| No | 90 | 91.8% |
| How did you think the policy change was achieved (n = 14) |  |  |
| Notified Hospital Management | 3 | 21.4% |
| Collaborated with policy makers | 3 | 21.4% |
| The findings is cited globally/Published | 2 | 14.3% |
| Included in the management protocol | 2 | 14.3% |
| Intervention Programme | 1 | 7.15% |
| Personal implementation | 1 | 7.15% |
| Liaised with WHO | 1 | 7.15% |
| Interaction and notification of relevant bodies | 1 | 7.15% |

\*\*: Some respondents were PI in more than one type of research. ‡ Some respondents gave more than one reasons for embarking on a research;

The list of challenges in descending order starting with the most challenging were getting sponsor 86 (78.2%), developing research proposal 80 (72.7%), and choosing research areas/topic 79 (71.8%) (Table3).

**Table 3:** Challenges in conduct of research

| Variables | No challenge<br>n (%) | Challengin<br>g (%) | Average<br>rating where<br>1 is the least<br>and 7 highest | 1<br>(Low<br>challenge) | 2 - 4<br>Medium<br>challenge) | 5 - 7<br>High<br>Challenge) |
| --- | --- | --- | --- | --- | --- | --- |
| Choosing Research Area/Topic | 31 (28.2) | 79 (71.8) | 2.26 (7<sup>th</sup>) | 34 | 36 | 9 |
| Developing Research Proposal | 30 (27.3) | 80 (72.7) | 2.60 (5<sup>th</sup>) | 22 | 50 | 8 |
| Getting Sponsor/Fund | 24 (21.8) | 86 (78.2) | 4.24 (1<sup>st</sup>) | 26 | 25 | 35 |
| Coordinating Field Work | 38 (34.5) | 72 (65.5) | 2.81 (3<sup>rd</sup>) | 19 | 41 | 12 |
| Analysis | 34 (32.7) | 74 (67.3) | 2.65 (4<sup>th</sup>) | 15 | 49 | 8 |
| Publishing of research findings | 50 (45.5) | 60 (54.5) | 3.43 (2<sup>nd</sup>) | 13 | 34 | 14 |
| Getting collaborators | 52 (47.3) | 58 (52.7) | 2.38 (6<sup>th</sup>) | 26 | 20 | 12 |
| Others (n= 9)<br>Default of subjects (n=3)<br>Parents/caregivers (n=3)<br>Time management (n=2)<br>Retirement of Grants (n= 1) |  |  |  |  |  |  |

**Figure 4:**
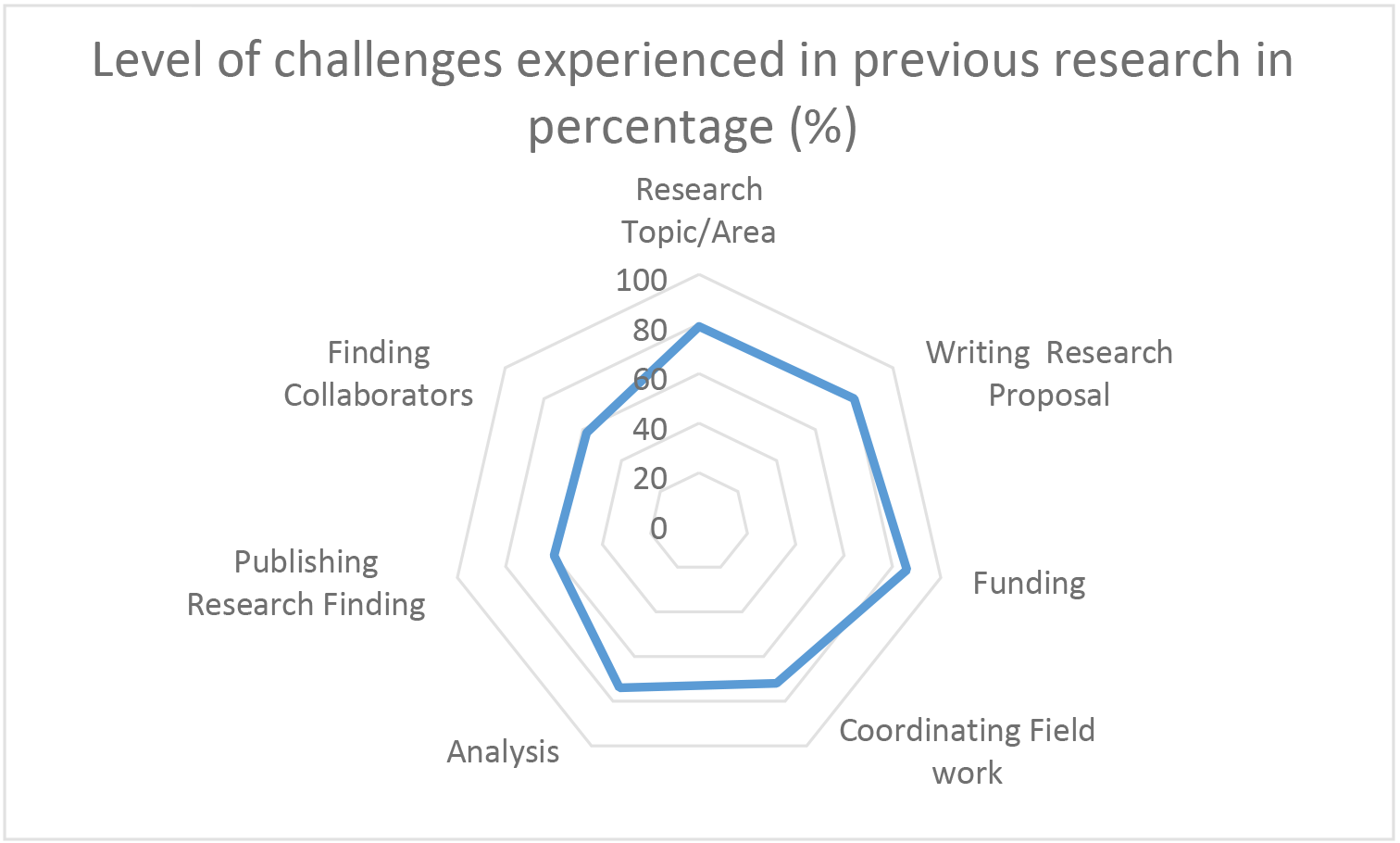
Level of challenges experienced in different components of research

To determine area that will require capacity development, respondents were asked to enlist research related trainings they had received. The least received trainings were data quality assurance (7.3%), and team science (8.2%) (Table 4).

**Table 4:** Aspects of Research the respondents had received training

| Aspect received training | N | % |
| --- | --- | --- |
| Research Ethics | 78 | 70.9 |
| Good Clinical Practice | 51 | 46.4 |
| Good Laboratory Practice | 17 | 15.5 |
| Responsible Conduct of Research | 40 | 36.4 |
| Crises Communication | 11 | 10 |
| Grant Writing | 29 | 26.4 |
| Manuscript Development | 39 | 35.5 |
| Biostatistics | 38 | 34.5 |
| Data Management | 35 | 31.8 |
| Data Quality Assurance | 8 | 7.3 |
| Research Monitoring | 12 | 10.9 |
| Monitoring and Evaluation | 22 | 20 |
| Budgeting | 13 | 11.8 |
| Team Sciences | 9 | 8.2 |

## Discussion

The majority of the research works carried out by paediatricians were prevalence studies. Although the challenges of conducting experimental studies involving children are obvious, the issues of funding and consenting process are outstanding. Clinical trials involving children are more challenging to carry out due to lack of funding, unique nature of children and ethical concerns. However current regulations and initiatives are improving the scope, quantity and quality of trials in children.^7^ Generous funding will dramatically increase the number of paediatricians participating in clinical experimental research. Presently, there are few experimental drug studies that involved children compared to adults.^8–10^ Part of the low level of involvement of children in research can be blamed on the existing guidelines which allow children to participate in experimental research only when their parents/caregivers give their consent. It is not surprising that only three participants in the study mentioned availability of subjects and issue of consent as challenges. The study shows low capacity of respondents to conduct high quality and more complex research. The situation may change when capacity is improved and more experimental research is conducted among paediatric subjects in Nigeria.^11^

The Belmont Report justifies inclusion of children in research. ^12, 13^ Interestingly among the tested capacity areas of research, research ethics and Good Clinical Practice (GCP) were the two instructional activities for which more paediatricians had received training. These trainings place paediatricians at vintage position to lead or be part of experimental study involving children.

The reported high prevalence of lack of access to grants is a disturbing factor to be urgently addressed failure of which will militate against conduct of high quality research targeted at improving the health children and adolescents in Nigeria. In accordance with the Convention on the Rights of the Child, ^14^ children have right to the highest attainable level of health care. This cannot be realized if children are not provided with health care based on proven evidence. Such evidence cannot be generated through extrapolation of researches outcomes in adults.^15–19^ The importance of this is highlighted by our finding that paediatricians were engaged in low quality research that does not feed into policy updates.

Evidence have shown that the expected outcome of any research work is to generate evidence that will change policy and practice.^20–23^ Unfortunately this study revealed that most of the studies carried out by paediatricians have not fulfilled this role. It appears that previous research efforts were not targeted at needs of children and adolescent. The driving force for research in the Nigeria includes academic pursuit and promotion for the academic paediatricians. The country must now move on to tailor research to the needs of children and adolescents. The real needs of children and adolescents have to be clearly identified and targeted by research. This way paediatricians will be exposed to the concept of targeted research for evidence that will inform policy improvement and practice (GRIPP).^24^ To achieve this goal there are basically two broad strategies to be employed: engaging the stakeholders is one and using evidence in decision making is the another.^21^ The sole goal of GRIPP is to achieve knowledge translation, knowledge transfer, knowledge exchange, research utilization, implementation, diffusion, and dissemination.^25^ The translation of research findings into actionable policy and programmatic guidance is an achievable goal to be exploited.

The study has shown that there is lack of strategic research capacity building in the country. It has always remained a point of argument; can clinicians be effective researchers? Beyond an obvious yes, a conscious and adequate capacity development is urgently needed. The capacity development should be ongoing through an organized structure that will enable and support programmes that will provide paediatricians irrespective of geographical place and areas of work with opportunities to interact and grow in their ability to plan and conduct clinical research. There are multiple means towards achieving this research capacity development. Strengthening formal education and routine training programme during the residency and creating internship for research skill acquisition and mentoring are key ways to attain optimum standards. ^26^ Building research capacity is very important. When research capacity is limited, as revealed in this study, there is a gap in the production of contextually relevant research evidence for communal benefits and in the synthesis of research evidence to inform practice, programmes and policies.

Lack of funding showed up strongly as a rate limiting step to conduct research by the respondents. The research component of the residency programme in Nigeria is poorly funded. There is also limited national funding arrangement for academic paediatricians to compete for. In order to respond to these concerns, capacity building programmes should receive high priority in health budgets. This can be in the form of career trajectories; institutional infrastructure and mentorship for research; tailoring research capacity efforts to a broad range of clinical and consultation services, scholarship and the involvement of community and policy makers in the process.^27^

This study revealed low questionnaire retrieval rate from respondents in an internet-based methodology and conference survey compared to individualized questionnaire administration survey. Similar low response rate from research participants have been previously reported ^28,29^ Doctors are very busy, they give most of their time to patient care. The internet based survey ought to be easier, fast, lower cost, ability to accumulate very large volume of interviews in a short space of time and no interviewer effects. Despite these advantages, online survey is not optimized in research among the populations studied. Perhaps the study participants would have been encouraged to utilize online survey through creating awareness during conferences and use of incentives like winning phone credit or some megabits at the completion of the survey. Such incentivized method was demonstrated in a survey where at the completion of a survey, participants were entered into lottery for which they stood a chance to win cinema tickets. ^30^

Conclusion: This survey has revealed areas of gap amongst Pediatricians. The Nigerian Academic Paediatricians need to be stimulated to develop interest in research by building their presently low research capacity if the future paediatric practice is to be driven significantly by evidence. Paediatricians in Nigeria should be re-orientated to aggressively contribute to evidence-based practice by seeking to build their own capacity for high quality research and to demand strong support for research into children and adolescents’ health concerns in Nigeria. This will pave the way for programing of priority intervention for capacity building.

## References

1. NSW Health, Clinical Nurse Consultants-Domains and Functions. Department of Health North Sydney, Australia, 2005

2. Australian Government Department of Health and Ageing. Strategic Review of Health and Medical Research Final Report. Canberra: Commonwealth of Australia; 2013.)^*^

3. Wilkes Lesley, Cummings Joanne, Mckay Nicola. Developing a culture to facilitate research capacity buiding for clinical nurse consultants in generalist paediatric practice. Nursing research and Practice; 2013 http://dx.doi.org/10.1155/2013/709025)

4. K. Vaughan, L.M. Wilkes, J. O’Baugh, and R. O’Donohue, “The role and scope of the clinical nurse consultant in Wentworth area health service:a qualitative study,” Collegian.2005;12(3):14–19

5. Dana Burde, Joel Middleton, Cyrus Samii. Research Capacity Assessment Scoring Framework, 2016, available https://www.alseproject.com

6. Survey Methods. www.surveymethods.com. Accessed on 4th May 2018

7. P.D. Joseph, J.C. Craig, P.HY Caldwell. Clinical trials in children. BJCP 2015;79(3):357–369

8. Conroy S, Choonara I, Impicciatore P, et al. Survey of unlicensed and off label drug us in paediatric wards in European countries. BMJ 2000;320:79–82.

9. Choonara I. Clinical trials of medicines in children. BMJ 2000;321:1093–4.

10. Smyth RL. Research with children. BMJ 2001;322:1377–8.

11. D.S. Wendler Assent in paediatric research: theoretical and practical considerations. J Med Ethics 2006;32:229–234. Doi:10.1136/jme.2004.011114

12. The National Commission for the Protection of the Human Subjects of Biomedical and Behavioral Research. The Belmont Report: Ethical Principles and Guidelines for Protection of Human Subjects of Research. HHS.gov. US Department of Health and Human Services. [Accessed on 2018 May 5]. Available from: http://www.hhs.gov/ohrp/humansubject/guidance/belmont.html

13. S. B. Bavdekar. Pediatric clinical Trials. Perspect Clin res 2013;4(1):89–99.

14. General Assembly of the United Nations. Convention on the Rights of the Child, 20 November 1989. [Last accessed on 2018 April 19]. Available from: http://www.unicef.org/crc/crc.htm

15. Bavdekar SB, Sadawarte PA, Gogtay NJ, Jain SS, Jadhav S. Off-label drug use in a Pediatric Intensive Care Unit. Indian J Pediatr. 2009;76:1113–8

16. Jain SS, Bavdekar SB, Gogtay NJ, Sadawarte PA. Off-label drug use in children. Indian J Pediatr. 2008;75:1133–6.

17. Williams K, Thomson D, Seto I, Contopoulos-Ioannidis DG, Ioannidis JP, Curtis S, et al. Standard 6: Age Groups for Pediatric Trials. Pediatrics. 2012;129:S153–60

18. Dunne J, Rodriguez WJ, Murphy MD, Beasely BN, Burckart GJ, Filie JD, et al. Extrapolation of adult data and other data in pediatric drug-development rograms. Pediatrics. 2011;128:e1242–9.

19. Gill D, Kurz R. Practical and Ethical Issues in Pediatric Clinical Trials. Appl Clin Trials. 2003;12:41–4

20. Lavis JN, Oxman AD, Lewin S, Fretheim A. SUPPORT Tools for evidenceinformed health Policymaking (STP) 3: setting priorities for supporting evidence informed policymaking. Health Res Policy Syst. 2009;7:S3.

21. Mirzoev T, Green A, Gerein N, Pearson S, Bird P, Ha BT, Ramani K, Qian X, Yang X, Mukhopadhyay M, Soors W. Role of evidence in maternal health policy processes in Vietnam, India and China: findings from the HEPVIC project. Evid Policy. 2013;9:493–511.

22. Alliance for Health Policy and Systems Research. Strengthening health systems: the role and promise of policy and systems research. Geneva: WHO, Alliance for Health Policy and Systems Research; 2004.

23. Green A, Bennett S. Sound Choices: Enhancing Capacity for EvidenceInformed Health Policy. AHPSR Biennual Review. Geneva: WHO, Alliance for Health Policy and Systems Research; 2007

24. B. Uzochukwu, O. Onwujekwe, C. Mbachu, E. Etiaba, M.E. Nystrom, L. Gilson. The challenge of bridging the gap between researchers and policy makers: experiences of a Health Policy Research Group in engaging policy makers to support evidence informed policy making in Nigeria. Globalization and Health 2016;12:67.doi:10.1186/s12992-016-0209-1

25. Oxman AD, Lavis JN, Lewin S, Fretheim A. SUPPORT Tools for evidence-informed health Policymaking (STP) 1: What is evidence-informed policymaking? Health Res Policy Syst. 2009;7 Suppl 1:S1. doi:10.1186/1478-4505-7-S1-S1.

26. M. Murray, Beyond the myths and magic of mentoring: How to facilitate an effective mentoring process, Jossey-Bass, San Francisco, Carlif, USA, 2001

27. N. Edwards, D. Kaseje, E. Kahwa. In Building and Evaluating research capacity in Heathcare System: Case Studies and Innovative Models. Eds N. Edwards, D. Kaseje, E. Kahwa, UCT Pub 2016

28. B. Duffy, K. Smith, G. Terhanian, J. Bremer. Comparing data from online and face-to-face surveys. International Journal of Market Research 2005;47(6):615–639

29. G. Szolnoki, D. Hoffmann. Online, face-to-face and telephone surveys - comparing different sampling methods in wine consumer research. Wine Economis and Policy 2013;2(2):57–66

30. Marie L Misso, dragan Ilic, Terry P Haines, Alison M Hutchinson, Christine E East, Helena J Teede. Development, implementation and evaluation of a clinical research engagement and leadership capacity building program in a large Australian healthcare service. BMC Medical Education 2016;16:133 doi:10.1186/s12909-016-0525-4

